# Characterization of nuclear pore complex targeting domains in Pom152 in *Saccharomyces cerevisiae*

**DOI:** 10.1101/2020.06.18.157685

**Authors:** Jacqueline T. Brown, Alexandra J. Haraczy, Christopher M. Wilhelm, Kenneth D. Belanger

## Abstract

Pom152 is a transmembrane protein within the nuclear pore complex (NPC) of fungi that is important for NPC assembly and structure. Pom152 is comprised of a short amino-terminal region that remains on the cytosolic side of the nuclear envelope (NE) and interacts with NPC proteins, a transmembrane domain, and a large, glycosylated carboxy-terminal domain within the NE lumen that self-assembles to form the NPC membrane ring. Here we show that the N-terminal 200 amino acids of Pom152 that include only the amino-terminal and transmembrane regions of the protein are sufficient for localization to the NPC. Full-length, glycosylation-deficient, and truncated Pom152-GFP chimeras expressed in cells containing endogenous Pom152 localize to both NPCs and cortical endoplasmic reticulum (ER). Expression of Pom152-GFP fusions in cells lacking endogenous Pom152 results in detectable localization at only the NE by full-length and amino-terminal Pom152-GFP fusions, but continued retention at both the NE and ER for a chimera lacking just the carboxy-terminal 377 amino acids. Targeted mutations in the amino-terminal and transmembrane domains did not alter Pom152 localization and neither deletion of Pom152 nor its carboxy-terminal glycosylation sites altered the nuclear protein export rate of an Msn5/Kap142 protein cargo. These data narrow the Pom152 region sufficient for NPC localization and provide evidence that alterations in other domains may impact Pom152 targeting or affinity for the NPC.

## Introduction

Nuclear pore complexes (NPCs) are large, aqueous, proteinaceous channels that perforate the nuclear envelope and regulate communication and transport between the nucleoplasm and cytoplasm (Beck & Hurt, 2017; Knockenhauer & Schwartz, 2016; Tran & Wente, 2006). Each NPC is composed of about 30 different proteins termed nucleoporins (Nups), with each Nup present in 8 to 32 copies (Alber, Dokudovskaya, Veenhoff, Zhang, Kipper, Devos, Suprapto, Karni-Schmidt, Williams, Chait, Rout, et al., 2007; Kim et al., 2018; Mi et al., 2015; Rajoo et al., 2018). Nups can be categorized by grouping them based on their function and characteristic functional domains: FG-Nups, non-FG-Nups, and pore membrane Nups. FG-Nups facilitate substrate translocation across the NPC via hydrophobic regions containing phenylalanine-glycine (FG) repeat sequences by forming a diffusion barrier within the pore that selectively allows passage of nuclear transport factors (NTFs) and their cargoes (Onischenko & Weis, 2011; Rout et al., 2000). Non-FG Nups form the structural scaffold with which FG-Nups associate within the NPC (Kim et al., 2018; Kosinski et al., 2016; Lin et al., 2016). The pore membrane Nups (POMs) span the pore membrane, the lipid bilayer within the NPC that connects the inner and outer nuclear membranes. In yeast, the pore membrane Nups are Pom152, Pom34, and Ndc1, and each has a single transmembrane domain and is involved in both assembly of nascent NPCs and maintenance of the structural organization of each NPC via association with each other and with non-FG Nups within the pore (Kim et al., 2018; Onischenko et al., 2009).

Pom152 is a typeII transmembrane protein with an amino-terminal cytosolic domain, a transmembrane (TM) domain spanning residues 176 – 195, and a carboxy-terminal lumenal domain that comprises most of the mass of the 1337 amino acid polypeptide (Tcheperegine et al., 1999; Wozniak et al., 1994). The carboxy-terminal domain self-oligomerizes within the NE lumen to generate the eight-fold symmetry of the NPC and, together with Pom34 and Ndc1, forms the membrane ring complex at the core of each NPC (Kim et al., 2018; Upla et al., 2017; Yewdell et al., 2011). The C-terminal domain contains at least four N-linked glycosylation sites, although the function of this glycosylation remains unclear (Belanger et al., 2005; Wozniak et al., 1994; Yewdell et al., 2011). The amino-terminal, cytosolic Pom152 domain associates with non-FG Nups to form the scaffold around the equator of the NPC that provides the structure on which the FG-Nups assemble (Alber, Dokudovskaya, Veenhoff, Zhang, Kipper, Devos, Suprapto, Karni-Schmidt, Williams, Chait, Sali, et al., 2007; Kim et al., 2018). Evidence suggests that Pom152 is present in either eight ((Mi et al., 2015; Rajoo et al., 2018) or 16 (Alber, Dokudovskaya, Veenhoff, Zhang, Kipper, Devos, Suprapto, Karni-Schmidt, Williams, Chait, Rout, et al., 2007; Kim et al., 2018; Rout et al., 2000) copies per NPC.

Here we further examine the Pom152 sorting determinant responsible for localization at the yeast NPC and investigate the role of Pom152 in NPC function. We observe that the amino-terminal 200 amino acids of Pom152 are sufficient for targeting to NPCs. Expression of GFP-tagged versions of Pom152 result in both nuclear envelope and cortical endoplasmic reticulum localization in cells expressing endogenous Pom152. Eliminating the expression of wild type Pom152 decreases cortical ER localization of all Pom152-GFP chimeras tested except for a truncation lacking 377 amino acids from the carboxy-terminus. Pom152 mutants completely lacking the carboxy-terminus in conjunction with mutations of select residues within or adjacent to the transmembrane sequence showed localization comparable to wild type Pom152, suggesting that at least these portions of the amino-terminal and TM domains are not required for Pom152 direction to the NPC. We also observe that deletion of *POM152* or removal of its lumenal glycosylation sites does not significantly alter the kinetics of Msn5-mediated nuclear protein export from yeast nuclei, consistent with observations that *pom152* mutants do not have nuclear protein import or mRNA export defects (Belanger et al., 2005; Madrid et al., 2006). These data add to our understanding of Pom152 domain structure and function within the NPC and targeting to NPCs.

## Materials and Methods

### Yeast strains, plasmids, and molecular techniques

Yeast strains (Table 1) and plasmids (Table 2) used in this study are listed in the tables below. Introduction of DNA into yeast was performed by lithium acetate transformation (Gietz & Woods, 2006) and growth and selection were performed using established protocols. *POM152-GFP* (pKBB463), *pom152Δglyc-GFP* (pKBB464), pom152^1-960^-GFP (pKBB521), and pom152^1-^200-GFP (pKBB520) fusions were generated by amplifying sGFP from pJK19-1 using PCR with PfuUltra™ HF (Agilent, Inc. Santa Clara, CA) and primers that generated chimeric Pom152-sGFP PCR products for homologous recombination in yeast. For pKBB463 and pKBB464, primers KOL236 (5’-GAAATTACAGATGCTTATTGTTTTGCCAAAAATGATCTTTTTTTCAATAACGCTAGCAAAGGAGAAGAACTC-3’) and KOL307 (5’-ATTTCTGTGGATGTTCAAAAGTCTGCTTTTAACACACCTCTATAGACCGTCCTTTCGGGCTTTGTTAGCAGCC-3’) were used for amplification and the resulting PCR products were cotransformed into yeast strain BY4742 with either plasmid pPM1-HA (Wozniak et al., 1994) and pKBB382 (Belanger et al., 2005) linearized with restriction endonuclease *Psh*AI. Primers used to amplify sGFP from pJK19-1 were KOL407 (5’-AGAAGAAGTCTGTCAAGGGATGGAAGGTACGGTTGATTTGGCTCTATTTGGTTCTCCAGCTAGCAAAGGAGAAGAACTC-3’) and KOL307 for pKBB521 and KOL406 (5’-CAGATTTTAGCTATGCTACTATTGAACATTTTCATATCAAGCGATCACGAGTTCGTTGCTAGCAAAGGAGAAGAACTC-3’) and KOL307 for pKBB520, and PCR products were independently cotransformed into BY4742 with *Psh*AI-linearized pPM1-HA. Cells containing recombinant plasmids were selected for on SD-Leu media and Pom152-sGFP fusions were confirmed by PCR and DNA sequencing. Site-directed mutagenesis with a Stratagene QuikChange Lightning Multi Site-Directed Mutagenesis Kit (Agilent, Inc. Santa Clara, CA) was used to generate *pom152^Y180L,Q182L^-GFP* (pKBB553) and *pom152^S158A,F159A,F160A^-GFP* (pKBB556) from *pom152^1-200^-GFP* (pKBB520) using mutagenic oligonucleotides KOL434 (5’-GCCATGGGTTGTTTTACTCCTGATTTTAGCTATGC-3’) and KOL437 (5’-GTATTTCATTATAGATGCCGCCGCCCTGTATGTTTTACCATCC-3’), respectively, as per manufacturer instructions. Successful mutagenesis was confirmed by DNA sequencing.

**Table 1.** Yeast strains used in this study.

| **Strain** | **Genotype** | **Source** |
| --- | --- | --- |
| BY4742 | *MATα his3 leu2 lys2 ura3* | Open Biosystems |
| KBY776 | *MATα pom152ΔKAN<sup>R</sup> his3 leu2 lys2 ura3* | Open Biosystems |
| KBY850 | *MATα nup133ΔKAN<sup>R</sup> his3 leu2 lys2 ura3* | Open Biosystems |
| KBY893 | *MATα msn5ΔKAN<sup>R</sup> his3 leu2 lys2 ura3* | Open Biosystems |
| KBY1292 | *MATα nup120ΔKAN<sup>R</sup> his3 leu2 ura3 lys2* | Open Biosystems |
| Y5563 | *MATα can1Δ::MFA1pr-HIS3 lyp1 leu2 met15 ura3 his3* | (Tong & Boone, 2006) |
| KBY1377 | *MATα can1Δ::MFA1pr-HIS3 lyp1 leu2 met15 ura3 his3*<br>*pom152Δglyc-NAT<sup>R</sup>* | This study |
| KBY1378 | *MATα can1Δ::MFA1pr-HIS3 lyp1 leu2 met15 ura3 his3*<br>*pom152ΔNAT<sup>R</sup>* | This study |

**Table 2.** Plasmids used in this study.

| **Plasmid name** | **Genotype** | **Source** |
| --- | --- | --- |
| pRS315 | *CEN LEU2 Amp<sup>R</sup>* | (Sikorski & Hieter, 1989) |
| pJK19-1 | pET12a-sGFP | Pamela Silver |
| pLMB127 | *CEN URA3 3x-GFP-Crz1 Amp<sup>R</sup>* | (Boustany & Cyert, 2002) |
| pKW1803 | YIP-Lac *dsRed-HDEL NatMX* | (Madrid et al., 2006) |
| pPM1-HA | *CEN LEU2 Amp<sup>R</sup> POM152-HA* | (Wozniak et al., 1994) |
| pKBB382 | *CEN LEU2 Amp<sup>R</sup> pom152Δglyc-HA* | (Belanger et al., 2005) |
| pKBB463 | *CEN LEU2 Amp<sup>R</sup> POM152-HA-sGFP* | This study |
| pKBB464 | *CEN LEU2 Amp<sup>R</sup> pom152Δglyc-sGFP* | This study |
| pKBB520 | *CEN LEU2 Amp<sup>R</sup> pom152<sup>1-200</sup>-sGFP* | This study |
| pKBB521 | *CEN LEU2 Amp<sup>R</sup> pom152<sup>1-960</sup>-sGFP* | This study |
| pKBB553 | *CEN LEU2 Amp<sup>R</sup> pom152<sup>Y180L,Q182L</sup>-sGFP* | This study |
| pKBB556 | *CEN LEU2 Amp<sup>R</sup> pom152<sup>S158A,F159A,F160A</sup>-sGFP* | This study |

### Fluorescence Microscopy

Yeast strains (Table 1) were co-transformed with plasmids expressing alleles of Pom152-GFP (Table 2) and pKW1803 (HDEL-dsRed) linearized with *Eco*RV (Madrid et al., 2006). For fluorescence microscopy, cells were grown overnight at 24°C in SD–Leu media to early log phase (A_600_ = 0.05–0.2) and visualized under direct fluorescence using a Nikon E400 microscope. KBY850 (*nup133Δ*) strains were shifted to 35°C for 1 hour prior to microscopy. Images were captured using an RTKE Spot CCD camera and SPOT-RTKe Imaging software (Diagnostic Instruments, Inc).

### Crz1 nuclear protein export assays

Kinetic analyses of Crz1-GFP nuclear export in wild type and *POM152* mutant strains were performed as described previously (Finn et al., 2013). Briefly, pLMB127 (Boustany & Cyert, 2002) was transformed into yeast strains BY4742, KBY776, KBY893, KBY1377, and KBY1378 to allow for Crz1-GFP expression. Crz1-GFP was induced to accumulate in the nucleus by inclusion of 170 mM CaCl_2_ in SD-Leu media, and Crz1-GFP nuclear export was initiated by the addition of .5 μg/ml FK506. The percent of cells exhibiting distinctly nuclear GFP fluorescence was calculated by direct observation immediately before FK506 addition (t=0) and at two minute intervals after. Export assays were performed blind with each yeast strain assayed at least three times and greater than 100 cells observed at each timepoint.

## Results

### The amino-terminal and transmembrane domains together are sufficient for localization of Pom152 to the yeast NPC

In order to further define the region of Pom152 sufficient for anchoring at the NPC, Pom152-GFP chimeras containing either wild type (wt) *POM152* (Pom152-GFP), the amino-terminal and TM sequences exclusively (residues 1-200; pom152^1-200^-GFP), a truncation of the carboxy-terminal 377 amino acids (residues 1-960; pom152^1-960^-GFP), or *POM152* lacking glycosylation sites (pom152^Δglyc^-GFP) were transformed into wild type yeast (Fig. 1A). These Pom152-GFP-expressing plasmids were co-transformed with a plasmid encoding the endoplasmic reticulum (ER) targeting sequence HDEL fused to the red fluorescent protein DsRed (HDEL-DsRed, (Madrid et al., 2006)) and observed by phase contrast and fluorescence microscopy. When expressed in wild type yeast, all Pom152-GFP chimeras localized to the NE and likely to the NPCs as evidenced by the characteristic punctate GFP-fluorescence pattern around the nuclear rim (Fig. 1B. left panel). Each Pom152 fusion also exhibited some co-localization with HDEL-DsRed fluorescence, indicating targeting to or retention within the cortical ER. To test if intracellular localization of the Pom152-GFP chimeras was affected by the presence of endogenous wt Pom152, we co-transformed plasmids expressing the same four Pom152-GFP chimeras and the HDEL-DsRed ER marker into yeast lacking *POM152* (*pom152Δ*) and observed transformed cells by phase contrast and fluorescence microscopy. Localization of Pom152^wt^, *pom152*^Δglyc^ and *pom152*^1-200^ to the cortical ER was reduced to the point of no longer being detectable, but *pom152*^1-960^ retained significant localization to the cortical ER (Fig. 1B, right panel). These observations suggest that no portion of the lumenal carboxy-terminus of Pom152 is required for NPC localization, but that alterations in the carboxy-terminal domain may alter NPC affinity or targeting relative to wild type.

**Figure 1.**
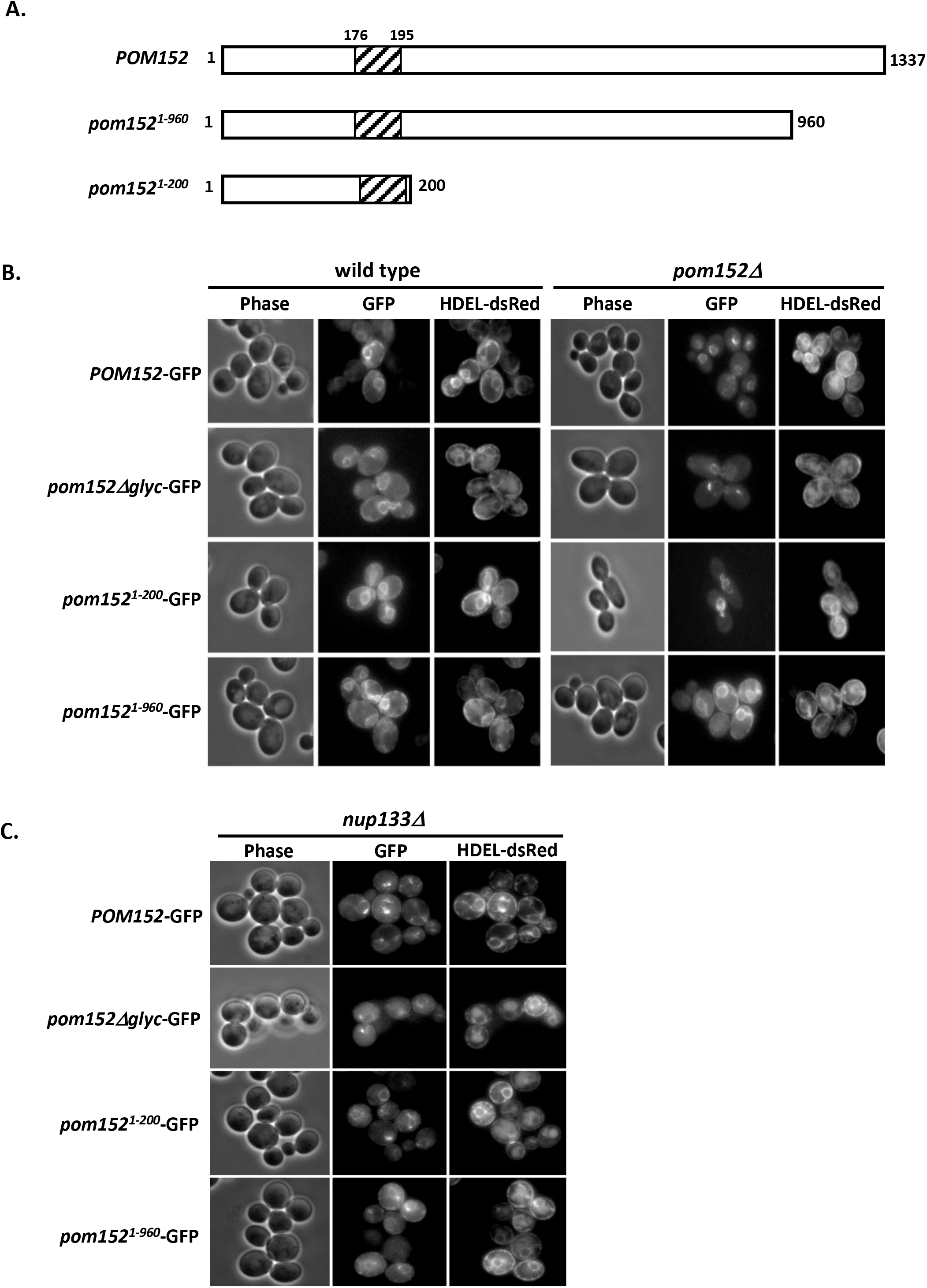
Removal or deglycosylation of the Pom152 carboxy-terminal, lumenal domain does not alter localization at the NPCs, while truncation of a portion of the lumenal domain results in increased ER localization. **(A)** Cartoon schematic of Pom152 carboxy-terminal truncations containing the first 960 (*pom152^1-960^*) or 200 (*pom152^1-200^*) amino acids of Pom152 expressed as chimeras with GFP. **(B)** Wild type *POM152-GFP*, a mutant lacking four glycosylation sites from the carboxy-terminal, lumenal domain (*pom152Δglyc-GFP*), and C-terminal truncations *pom152^1-200^-GFP* and *pom152^1-960^-GFP* were expressed in wild type yeast and yeast lacking endogenous *POM152* (*pom152Δ*). Cells were co-transformed with a plasmid expressing the endoplasmic reticulum marker HDEL-dsRed (Madrid et al. 2006) and observed via phase-contrast and fluorescence microscopy. *pom152-GFP* mutants localize to both the nuclear envelope and ER in cells expressing endogenous *POM152*, while only the *Pom152^1-960^-GFP* retains ER localization in *pom152Δ* cells. **(C)** Expression of *Pom152-, pom152Δglyc-, pom152^1-^ 200-* and *pom152^1-960^-GFP* fusions in cells lacking Nup133 (*nup133Δ*) results in clustered foci of fluorescence indicating NPC localization of all Pom152-GFP chimeras tested.

The partial localization of *pom152*^1-960^ to the endoplasmic reticulum in both wt and *pom152*Δ yeast raised the possibility that the perceived NPC localization of the Pom152-GFP fusions might be due to localization to the nuclear envelope as a consequence of this membranous structure being contiguous with the cortical ER. To confirm NPC localization, we expressed the same four Pom152-GFP chimeras in *nup133*Δ cells that exhibit a temperature-sensitive, NPC clustering phenotype (Belgareh & Doye, 1997) and contain the HDEL-dsRed ER marker. Phase contrast and fluorescence microscopy after incubation at the restrictive temperature showed all four Pom152-GFP fusions exhibiting clustered fluorescence around one side of the nuclear rim, characteristic of NPC localization in *nup133Δ* cells (Fig. 1C). A small amount of each Pom152-GFP remained in the cortical endoplasmic reticulum around the periphery of the cell. These observations confirm the localization of these Pom152-GFP chimeras to NPCs and that the first 200 amino acids of Pom152 are sufficient for NPC localization.

### Alteration of specific residues within and adjacent to the Pom152 transmembrane sequence does not affect localization to the yeast NPC

Metazoan NPCs contain a single-pass transmembrane protein named gp210/Nup210 that exhibits some similarities to Pom152, including involvement in NPCs assembly and structural organization (Stavru et al., 2006). Nup210 also has a large lumenal domain and a smaller cytosolic region like Pom152, but the topology of the protein is switched such that the N-terminus is in the NE lumen and the C-terminus is cytosolic (Greber et al., 1990). Interestingly, the transmembrane domain alone of Nup210 is sufficient for targeting to the NPC, as is a 20 amino acid region of the cytosolic domain of the protein (Wozniak & Blobel, 1992). Because the Nup210 transmembrane sequence and cytosolic domain each contain sorting determinants sufficient for localization to the mammalian NPC (Wozniak & Blobel, 1992), we sought to identify sequences that function similarly in and adjacent to the Pom152 transmembrane and cytosolic domains. Although the amino acid sequence of Pom152 does not align with Nup210 in this region, a query of BLAST (Altschul et al., 1990) using the first 200 amino acid residues from Pom152 reveals significant homology across fungal organisms, including some residues that are conserved across nearly all species examined (Fig. 2A). Among these conserved residues are a serine at residue 158 (S158) in *S. cerevisiae* Pom152, a phenylalanine at residue 160 (F160), a glutamine at residue 182 (Q182). We used site-directed mutagenesis to create two mutant *pom152*^1-200^-GFP fusions (Fig. 2B): one in which Tyr180 and Gln182 are both replaced by Leu (*pom152*^Y180L,Q182L^-GFP) and another in which Ser158-Phe159-Phe160 are changed to Ala residues (*pom152*^S158A,F159A,F160A^-GFP). Yeast strains deleted for *POM152* (*pom152*Δ, Fig. 2C) expressing each transmembrane domain mutant and HDEL-dsRed were observed by phase contrast and fluorescence microscopy. Both *pom152* transmembrane mutants localized to the NPC with no observable difference from wild type Pom152-GFP or the pom152^1-200^-GFP truncation (compare Figs. 1B and 2C), suggesting that these specific residues of the TM sequence or its flanking cytosolic sequence are not required for Pom152 targeting to or anchoring at the pore membrane domain.

**Figure 2.**
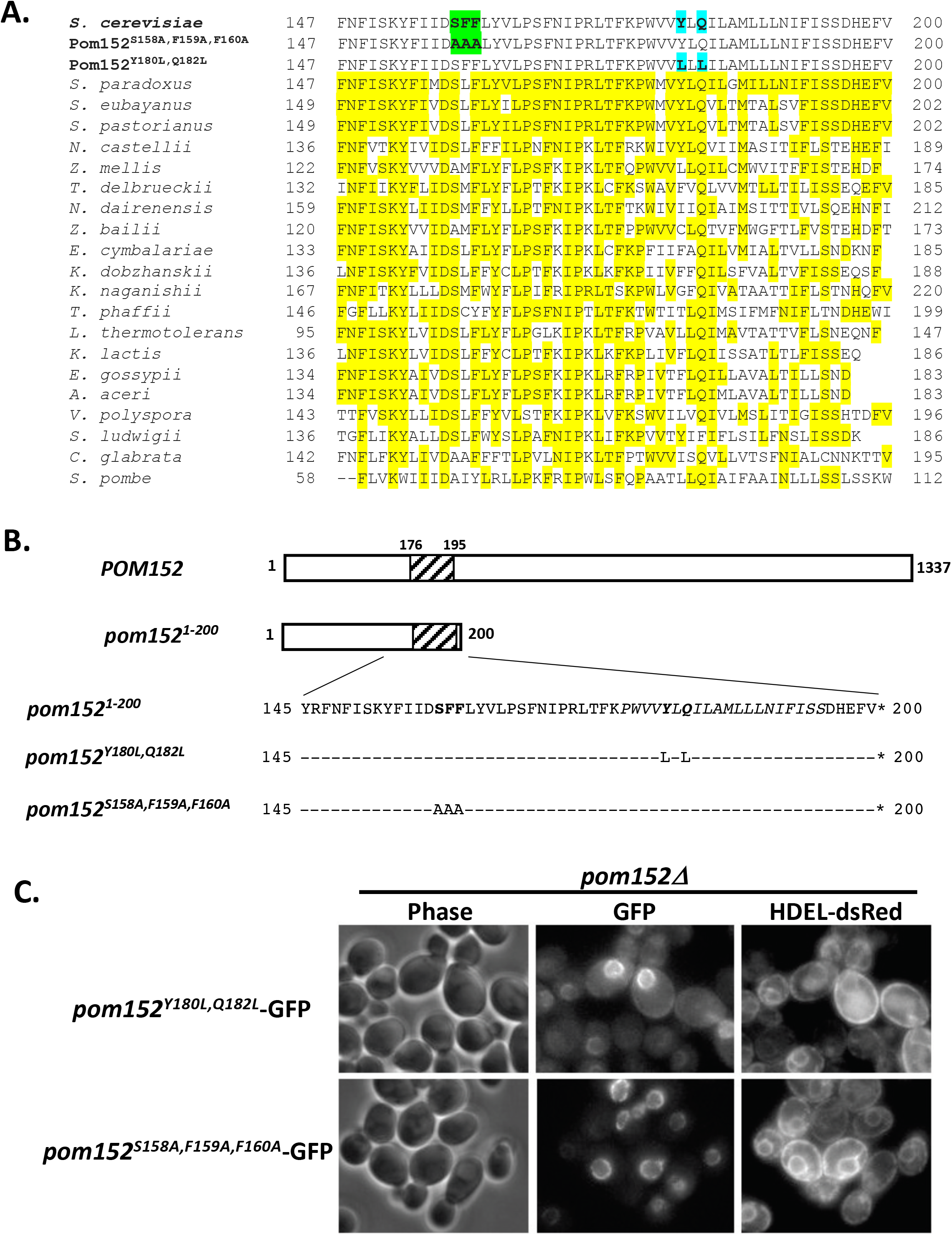
Targeted mutations within or flanking the transmembrane domain of Pom152 do not affect localization to the nuclear envelope. **(A)** Alignment of amino acid residues 147 – 200 from *S. cerevisiae* Pom152 with conserved regions of Pom152 from other fungi. Yellow highlights indicate identities with *S. cerevisiae*. Green highlights depict residues altered in *pom152^S158A,F159A,F160A^-GFP* and blue highlights depict those mutagenized in *pom152^Y180L,Q182L^-GFP.* **(B)** Schematic of Pom152 regions used to make *POM152-GFP* and *pom152^1-200^-GFP, pom152^Y180L,Q182L^-GFP* and *pom152^S158A,F159A,F160A^-GFP* chimeras. **(C)** Phase-contrast (phase) and fluorescence imaging of yeast deleted for *POM152* (*pom152Δ*) co-expressing *pom152^1-200^-GFP* mutants with the ER marker HDEL-dsRed. The *pom152^Y180L,Q182L^-GFP* and *pom152^S158A,F159A,F160A^-GFP* fusions both localize to the nuclear envelope.

### Pom152 and Pom152 glycosylation is not essential for Crz1 nuclear protein export

Deletions of *POM152* or elimination of four glycosylation sites from the Pom152 lumenal domain do not detectably alter nuclear protein import kinetics or mRNA nuclear export (Belanger et al., 2005; Madrid et al., 2006). To examine whether a loss of Pom152 or its glycosylation altered the rate of facilitated nuclear export of a protein that had accumulated in the nucleus, we performed kinetic assays for Msn5/Kap142-mediated nuclear protein export on cells that had accumulated the transcription factor Crz1 in the nucleus and were subsequently induced to export the protein to the cytoplasm. Live-cell fluorescence microscopy showed rapid export of Crz1 from yeast nuclei with a disappearance of detectable nuclear Crz1-GFP over a similar time span in wild type, *pom152Δ*, and *pom152Δglyc* cells (Fig. 3), indicating that neither a full deletion of *POM152* nor removal of the lumenal glycosylation sites significantly alters Msn5-mediated nuclear protein export.

**Figure 3.**
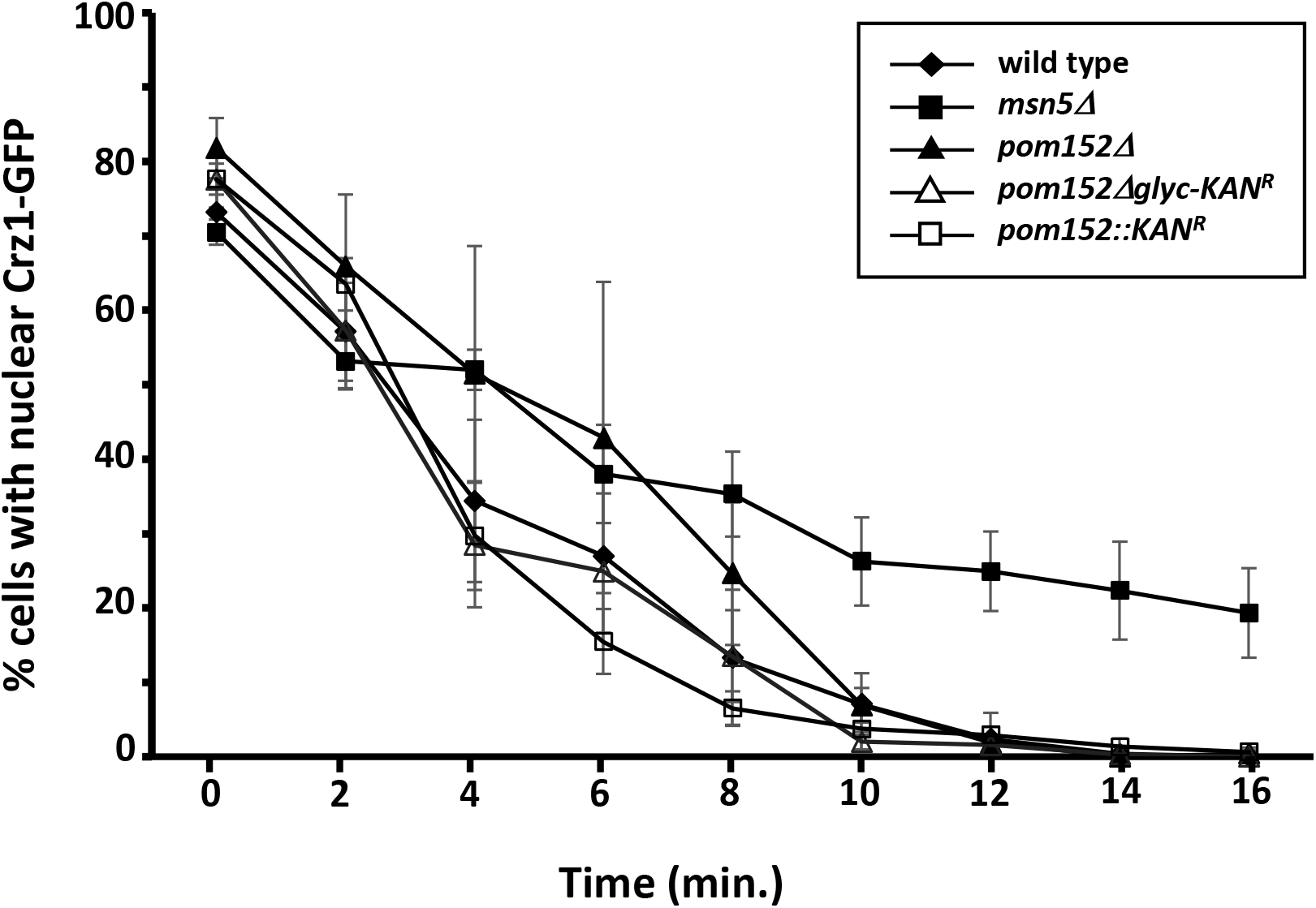
Pom152 deletion or removal of glycosylation sites does not alter Msn5-mediated nuclear protein export. Wild-type, *pom152Δ*, and *pom152Δglyc* cells expressing Crz1-GFP were allowed to accumulate Crz1-GFP in the nucleus in the presence of calcium and the percentage of cells with distinct nuclear fluorescence was recorded. Crz1 nuclear export was then induced by the addition of FK506. Data points indicate the percentage of cells with nuclear fluorescence for each strain. Error bars represent standard error of the mean.

## Discussion

In this study, we have presented evidence that amino-terminal amino acid residues 1-200 of the pore membrane nucleoporin Pom152, including the cytosolic and transmembrane regions, are sufficient for targeting the protein to the nuclear pore complex in *Saccharomyces cerevisiae*. Carboxy-terminal residues 201-1337 of Pom152 are not necessary for NPC localization, but removal of a portion of the carboxy-terminal domain may affect Pom152 targeting to and/or affinity for the NPC, as evidenced by continued cortical ER localization of pom152^1-960^-GFP, even after deletion of endogenous Pom152. Deletion of *POM152* or removal of its carboxy-terminal glycosylation sites do not alter the rate of nuclear export of the shuttling transcription factor Crz1.

### Pom152-GFP chimeras localize to both NPCs and cortical ER in the presence of endogenous Pom152

All of the Pom152-GFP chimeric proteins tested localize to both the NE and the cortical ER in wild type yeast expressing endogenous *POM152*. These data may suggest that co-expression of the recombinant Pom152-GFP fusions and endogenous Pom152 results in saturation of Pom152 sites within the NPC and that excess Pom152 not integrated into the nuclear pores is retained throughout the ER membrane. Alternatively, the observation of some Pom152-GFP in the ER membrane may indicate a lower affinity of the recombinant protein than endogenous Pom152 for the NPC, although the presence of some recombinant protein at the NE indicates that any reduction in affinity is not so great that Pom152-GFP chimeras are essentially excluded from NPCs. The observation that expression of Pom152-GFP, pom152Δglyc-GFP and pom152^1-200^-

GFP in cells lacking endogenous Pom152 results in NE envelope localization but undetectable fluorescence in the ER suggests that the affinity of these recombinant proteins for the NPC is sufficiently high for efficient targeting and/or retention. It has been shown previously that the amino-terminal region of Pom152 binds directly to the pore membrane Nups Pom34 and Ndc1 and to the non-FG nucleoporin Nup170 to link the membrane ring complex at the pore membrane to the inner ring complex that provides the inner scaffolding of the NPC (Kim et al., 2018; Makio et al., 2009; Onischenko et al., 2009) and this association likely mediates or stabilizes the Pom152 amino-terminal region at the NPC. Onischenko et al. (2009) observed that removal of the amino-terminal 170 amino acids of Pom152 alters NPC targeting, with the resulting truncation present at both the NPC and other intracellular membranes, likely predominantly cortical ER, suggesting that amino acids both within and downstream of the first 170 residues of Pom152 may be involved in NPC targeting and retention.

### Pom152^1-960^-GFP is present in cortical ER even in the absence of Pom152

Unlike the other Pom152-GFP proteins we tested, pom152^1-960^-GFP retains fluorescence in the ER as well as the NPC in *pom152*Δ yeast, suggesting that removal of the carboxy-terminal 377 amino acids from Pom152 may reduce NPC targeting or retention to the extent that a detectable fraction of the expressed protein is present in the ER. Alternatively, this truncated chimera may be more stable or may be overexpressed, resulting in excess pom152^1-960^-GFP accumulating in the ER. The carboxy-terminal region of Pom152 extends from residues 200 – 1337 and is composed of 8 or more immunoglobulin (Ig)-like folds which interact within the NE lumen and self-assemble head-to-tail and antiparallel to form the lumenal membrane ring at the equator of each NPC (Hao et al., 2018; Upla et al., 2017). Pom152^1-960^-GFP entirely lacks the carboxy-terminal three Ig-like folds and truncates a portion of the fourth. The Pom152 lumenal domain mediates Pom152 self-assembly within the membrane ring and all of the Ig-like folds are proposed to be involved in the anti-parallel association of juxtaposed Pom152 polypeptides (Hao et al., 2018; Kim et al., 2018; Upla et al., 2017; Yewdell et al., 2011). Thus, it is not unexpected that a loss of a portion of the Pom152 lumenal domain would alter NPC targeting of the protein.

### Mutations in the Pom152 transmembrane domain do not alter localization

Interestingly, the TM region alone of gp210/Nup210, the metazoan pore membrane NPC protein most structurally and functionally similar to Pom152, is sufficient for NPC localization (Wozniak & Blobel, 1992). Our limited mutagenesis of transmembrane (*pom152*^Y180L,Q182L^-GFP) and flanking amino-terminal (*pom152*^S158A,F159A,F160A^-GFP) residues in pom152^1-200^-GFP chimera did not reveal any changes in localization when expressed in wild type and *pom152Δ* cells and appeared to be targeted identically to the pattern observed for pom152^1-200^-GFP. Pom152 and gp210/Nup210 both are large, single-pass pore membrane nucleoporins with a large lumenal domain and shorter cytoplasmic region (Greber et al., 1990) that are important for NPC assembly and structural organization (Stavru et al., 2006). However, they contain only limited sequence similarity and are oriented such that the lumenal domain is at the carboxy-terminus of Pom152, but at the amino-terminus of gp210/Nup210 (Greber et al., 1990; Wozniak et al., 1994). However, yeast Pom152 expressed in mammalian cells localizes to NPCs (Wozniak et al., 1994), suggesting at least some conservation of intracellular targeting mechanism may occur. The development of a more comprehensive library of mutants and truncations will be necessary to more narrowly identify NPC targeting regions in Pom152 and determine if these sequences have any similarity to the gp210/Nup210 NPC targeting regions.

### Deletion of Pom152 does not alter Crz1 nuclear export kinetics

While previous work indicated that Pom152 alterations do not affect nuclear protein import and mRNA export through the NPCs (Belanger et al., 2005; Madrid et al., 2006), an analysis of the impact of Pom152 deletion or deglycosylation on nuclear protein export has not been reported. Here we show, not unexpectedly, that *pom152Δ* and *pom152Δglyc* mutants also do not exhibit significantly slowed rates of Msn5-mediated Crz1-GFP export from the nucleus. Despite the roles for Pom152 in NPC assembly and organization (Madrid et al., 2006; Marelli et al., 2001; Onischenko et al., 2009; Yewdell et al., 2011), direct and indirect physical associations with other nucleoporins in the NPC (Alber, Dokudovskaya, Veenhoff, Zhang, Kipper, Devos, Suprapto, Karni-Schmidt, Williams, Chait, Rout, et al., 2007; Alber, Dokudovskaya, Veenhoff, Zhang, Kipper, Devos, Suprapto, Karni-Schmidt, Williams, Chait, Sali, et al., 2007; Kim et al., 2018; Onischenko et al., 2009), and its synthetic genetic interactions with FG- and non-FG-*NUP* mutants (Belanger et al., 2005; Onischenko et al., 2009; Tcheperegine et al., 1999; Wozniak et al., 1994), to our knowledge deletions of *POM152* alone have yet to exhibit a phenotype relating to nuclear transport, NPC structure or distribution, or NPC assembly. This observation is likely a result of partial functional redundancy with the pore membrane Nups Ndc1 and Pom34, together with which Pom152 forms a complex in the pore membrane ring and interacts with non-FG Nups to organize the inner ring scaffold of the NPC (Alber, Dokudovskaya, Veenhoff, Zhang, Kipper, Devos, Suprapto, Karni-Schmidt, Williams, Chait, Sali, et al., 2007; Kim et al., 2018; Onischenko et al., 2009). Additional work is necessary to identify the exact role of Pom152 in NPC assembly and organization, the residues within Pom152 that mediate its targeting and function, and how a conserved protein performing these functions can be deleted without apparent nuclear transport, NPC structure, or cell growth or viability phenotypes.

## Conclusions

In this study, we have generated a series of alterations in the *Saccharomyces cerevisiae* pore membrane nucleoporin Pom152 and have used these recombinant proteins to examine Pom152 targeting to NPCs. We observe that a Pom152-GFP chimera entirely lacking the carboxy-terminal lumenal domain of the protein localizes to NPCs in a pattern indistinguishable from full length Pom152-GFP, suggesting that the targeting sequence is in the amino-terminal cytosolic and/or transmembrane regions. A shorter truncation of the lumenal domain increases retention in the cortical ER, raising the possibility that the self-assembling lumenal domain may also have a role in NPC targeting or retention. Pom152 alterations tested do not affect facilitated nuclear export of a shuttling transcription factor. In summary, the amino-terminal 200 amino acids of Pom152 target the protein to the NPCs in yeast.

## Acknowledgements

The authors thank Martha Cyert, Karsten Weis, Munira Basrai, Pamela Silver, and Richard Wozniak for their generous sharing of yeast strains and plasmids, and Karyn Belanger and Sue Geier for their expert technical contributions, organizational skills, and mentorship. We also thank undergraduate colleagues in the Belanger lab who provided valuable discussions and insights and especially acknowledge Caroline Adams, Nolan Sheppard, Sarah Kruse, Mohammed Rahman, Iustin Moga, and Elizabeth Rivers for their work on Pom152-related projects. This work was generously funded by Colgate University through the Faculty Research Council and the Stuart Updike Fund in support of undergraduate biology research.

## Notes

### Competing Interest Statement

The authors have declared no competing interest.

### Summary of Updates

Figure 2 has been revised to add an alignment with other fungal Pom152 sequences, and the text has been updated to include these data.

